# Comparative Transcriptomics of Mango (*Mangifera indica* L.) Cultivars Provide Insights of Biochemical Pathways Involved in Flavor and Color

**DOI:** 10.1101/196881

**Authors:** Waqasuddin Khan, Safina Abdul Razzak, M. Kamran Azim

## Abstract

Mango is an economically important fruit crop of many tropical and subtropical countries. Recently, leaf and fruit transcriptomes of mango cultivars grown in different geographical regions have characterized. Here, we presented comparative transcriptome analysis of four mango cultivars i.e. cv. *Langra*, cv. *Zill,* cv. *Shelly* and cv. *Kent* from Pakistan, China, Israel and Mexico respectively. De-novo sequence assembly generated 30,953-85,036 unigenes from RNASeq datasets of mango cultivars. KEGG pathway mapping of mango unigenes identified terpenoids, flavonoids and carotenoids biosynthetic pathways involved in flavor and color. The analysis revealed linalool as major monoterpenoid found in all cultivars studied whereas, monoterpene α-terpineol was specifically found in cv. *Shelly*. Ditepene gibberellin biosynthesis pathway was found in all cultivars whereas, homoterpene synthase involved in biosynthesis of 4,8,12-trimethyltrideca-1,3,7,11-tetraene (TMTT; an insect induced diterpene) was found in cv. *Kent*. Among sesquiterpenes and triterpenes, biosynthetic pathway of Germacrene-D, an antibacterial and anti-insecticidal metabolite was found in cv. *Zill* and cv. *Shelly*. Two bioactive triterpenes, lupeol and β-amyrin were found in cv. *Langra* and cv. *Zill*. Unigenes involved in biosynthesis of carotenoids, β-carotene and lycopene, were found in cultivars studied. Many unigenes involved in flavonoid biosynthesis were also found. Comparative transcriptomics revealed naringenin (an anti-inflammatory and antioxidant metabolite) as ‘central’ flavanone responsible for biosynthesis of an array of flavonoids. The present study provided insights on genetic resources responsible for flavor and color of mango fruit.

## Introduction

As a member of the family Anacardiaceae, mango (*Mangifera indica* Linn.) ranks second among tropical fruit crops after banana due to its rich sensational taste, color, aroma and huge economics significance (Srivastava et al. 2016; Litz 2009). Many mango varieties (i.e. cultivar abbreviated as cv.) are commercially grown in tropical and subtropical countries worldwide (Mukherjee and Litz 2009). According to Food and Agriculture organization of the United Nations (FAO), India holds the 1^st^ position in mango production followed by China, whereas Pakistan and Mexico rank 5^th^ and 6^th^ position respectively (FAOSTAT-2014; <u>www.faostat.fao.org</u>). Mango fruit is a rich source of bioactive phytochemicals including antioxidants and other health–promoting compounds (Lauricella et al. 2017; Fessard et al. 2017; Shah et al. 2010; Masibo and He 2009). This fruit is known for attractive colors, cherishing aroma, delightful taste and high nutritional value, due to its high content of vitamin C, carotenoids, flavones, terpenoids and minerals (Lauricella et al. 2017). The biochemical composition of mango pulp obtained from different mango cultivars varies with location of cultivation, variety, and stage of maturity (Dautt-Castro et al. 2015). Previous studies on mango were focused on the ripening process, volatile composition, antioxidant capacity, postharvest treatment and fruit quality (Srivastava et al. 2016; White et al. 2016; El-Hadi et al. 2013; Litz et al. 2009; Pino and Mesa 2006; Pino et al. 2005). Recently, Kuhn et al (2017) reported a consensus genetic map of mango using seven mapping population.

Recent transcriptome analysis of leaves and fruits of several mango cultivars by RNA– Seq provided insights of fundamental molecular biology of this plant. We first of all reported the mango leaf transcriptome of cv. *Langra* variety in 2014 (Azim et al. 2014). Mango fruit transcriptomes of cv. *Zill* (Wu et al. 2014), cv. *Shelly* (Luria et al. 2014), cv. *Kent* (Dautt-Castro et al. 2015), cv. *Dashehari* (Srivastava et al. 2016) and more recently cv. *Keitt* (Tafolla-Arellano et al. 2017) have been reported from China, Israel, Mexico, India and USA respectively. In another study, a leaf transcriptome identified genic-SSR markers and SNP heterozygosity in cv. *Armpali* (Mahoto et al. 2016). Simple sequence repeats (SSR) and SNP identification have proved to be informative DNA-based markers in plant molecular genetics (Mahoto et al. 2016; Khan and Azim 2011). Identification of SSR and SNP markers by transcriptome sequence datasets has potential to be utilized in mango breeding programs.

Genome-wide association, genetic mapping and identification of trait specific markers help to deploy important genes involved in flavor, aroma and pulp consistency for which mango is popularly consumed (Srivastava et al. 2016). Transcriptomic sequence datasets obtained from RNA-Seq of different mango cultivars resulted in 30,000–85,000 unigenes (Azim et al. 2014; Wu et al. 2014; Luria et al. 2014; Dautt-Castro et al. 2015; Srivastava et al. 2016). The transcriptome sequencing described in these reports, has been carried out using mango cultivars grown in different geographical regions. Hence, a comparative transcriptomic analysis was needed, in order to characterize common as well as cultivar and/or tissue-specific transcripts. Here, we present comparative analysis of transcriptomic datasets of available mango cultivars. This bioinformatics study identified genetic characteristics of mango responsible for its color, aroma, flavor and other agronomic traits at the systemic level.

## Materials and Methods

### Retrieval of Mango RNA-seq Data

For *de novo* assembly of RNA-Seq reads, the NGS reads of cv. *Langra* (SRA ID: SRR947746) (Azim et al. 2014), cv. *Zill* (SRA ID: SRP035450) (Wu et al. 2014), cv. *Shelly* (SRA ID: SRX375390) (Luria et al. 2014) and cv. *Kent* (SRA ID: SRP045880) (Dautt-Castro et al. 2015) were retrieved from NCBI Sequence Read Archive (SRA) (Leinonen et al. 2010). The cv. *Langra* sequence reads were from mango leaves where as other three were from mango fruits. In case of cv. *Shelly*, sequence reads of samples at different time intervals were pooled and used for subsequent analysis. For cv. *Kent*, sequence reads of mature mango RNA–Seq data were processed.

### Preprocessing of Mango RNA-seq Reads

Reads were converted from SRA format to fastq format using SRA Toolkit (<u>https://www.ncbi.nlm.nih.gov/sra/docs/toolkitsoft/</u>). Each SRA file provided two paired–end fastq files. The sequence reads were examined for quality by FASTQC (<u>https://www.bioinformatics.babraham.ac.uk/projects/fastqc/</u>) and were filtered by FASTX toolkit (<u>http://hannonlab.cshl.edu/fastx_toolkit/</u>) to obtain high–quality reads (reads with Q score ≥27). The processed forward (F1) and reverse (R2) read files were then paired using Pairfq script (<u>https://github.com/sestaton/Pairfq</u>). The headers of F1 and R2 files were configured according to CASAVA 1.8 format for Trinity (Grabherr et al. 2011) software using Fastool, a Trinity plugin.

### *De novo* Transcriptomic Assembly of Processed Reads

Transcriptome *de novo* assembly of clean reads was performed using Trinity (Grabherr et al. 2011) which uses three independent software modules – Inchworm, Chrysalis, and Butterfly – applied sequentially to process the sequencing data of RNA–seq reads. Bowtie aligner with some Perl scripts is part of Trinity pipeline. In brief, Trinity assembles the reads into unique sequences of transcripts, known as contigs. These contigs were clustered by constructing the complete *de Bruijn*graph for each cluster, and then partitioned the full read set among these disjoint graphs. Finally, Trinity processed the individual graphs in parallel, tracing the path that reads and pairs of reads take within the graph, ultimately reporting full–length transcripts for alternatively spliced isoforms, and teasing apart transcripts that corresponds to paralogous genes (Grabherret al. 2011). Consequently, we obtained four unigenes datasets corresponding to four mango cultivars.

### BLAST Analysis of Unigenes

Multiple BLAST strategies were used for sequence comparisons of unigenes against different sequences databases. (a) All unigenes from four assemblies were aligned against each other by BLASTN (E value cutoff ≤ 1e^−5^) to find the common and unique transcripts. The stringent alignment was defined by an E-value threshold of <1e^−10^, and a percent alignment and percent coverage length of ≥75%. (b) Unigenes in four datasets were aligned using BLASTN against the coding sequences (CDS) of *Citrus sinensis* (<u>https://www.citrusgenomedb.org/</u>)*, Citrus Clementina* (<u>https://www.citrusgenomedb.org/</u>), *Vitis Vinifera* (<u>http://www.plantgdb.org/VvGDB/</u>), *Ricinuscommunis* (<u>http://www.plantgdb.org/RcGDB/</u>) *and Populous tricocarpa* (<u>http://www.plantgdb.org/PtGDB/</u>) as well as against the plant–specific sequence (nucleotide and protein) databases of NCBI, UniProtKB, and Swiss–Prot with an Evalue cut–off ≤1e^−5^. (c) The assembled datasets were also BLASTed against NCBI non–redundant nucleotide (NT) and non–redundant (NR) protein sequence databases with E-value cut–off ≤1e^−5^.

### Functional Annotations of Unigenes

The assembled unigenes datasets were also filtered for redundant sequences using CD–HIT (Cluster Database at High Identity with Tolerance) (Fu et al. 2012) at 90% identity threshold. The non-redundant unigenes were further analyzed for coding regions using TransDecoder (<u>http://transdecoder.github.io/</u>). Obtained coding sequences (CDS) were then subjected to InterProScan v.5.15.54.0 (Jones et al. 2014) using IPRLOOKUP service for functional annotation and Gene Ontology (GO) assignments. Translated protein sequences were also scanned against the following InterPro signature databases: Hamap (201502.04), ProDom (2006.1), PRISF (3.01), SMART (6.2), TIGRFAM (15.0), PRINTS (42.0) and SUPERFAMILY (1.75). CateGOrizer (<u>http://www.animalgenome.org/tools/catego/</u>) was used to analyze GO term datasets into three GO classes’ i.e. biological process, cellular component and molecular function. InterProScan’s XML output along with their corresponding BLASTX result was loaded into Blast2GO java application (<u>https://www.blast2go.com/start–blast2go–2–8</u>) for gene annotations. The active biochemical pathway analysis was done by KAAS (Moriya et al. 2007) using KEGG database (Kanehisa 2002).

## Results

The transcriptomic *de novo* assembly resulted from Trinity (Grabherr et al. 2011) generated 30,953, 57,544, 58,797 and 85,036 of unigenes from cv. *Langra*, cv. *Zill*, cv. *Shelly* and cv. *Kent* mango cultivars. These unigenes were further characterized for functional annotations using BLAST and InterProScan (Jones et al. 2014). BLAST homology search showed 83-96% similarity of unigenes with sequences in Nt and Nr databases. The cultivar-specific consensus search among the four datasets of mango unigenes showed that 80–98% unigenes sequences were matched with each other with a similarity index of ≥75%. However, InterProScan identified 12,388, 29,303, 25,878, and 18,793 protein coding sequences in cv. *Langra*, cv. *Zill*, cv. *Shelly* and cv. *Kent* unigenes datasets respectively. Biochemical pathway analysis using KEGG-KASS (Kanehisa 2002; Moriya et al. 2007) identified numerous unigenes involved in the formation of genes that are involved for the production of biomolecules responsible for color and flavor of mango fruit.

## Discussion

Mango is an important fruit crop of many countries located in tropical and subtropical regions. In the absence of genome sequence information, transcriptomic sequences of different mango cultivars provided a wealth of data related to protein coding sequences. This study resulted in a ‘consensus transcriptome sequence reference’ obtained from four mango cultivars grown in Pakistan, China, Israel and Mexico. Initially, we retrieved 12.1, 68.4, 83.2, and 22.0 million paired-end RNA-Seq reads of cv. *Langra* (Pakistan), cv. *Zill* (China), cv. *Shelly* (Israel) and cv. *Kent* (Mexico) respectively from Sequence Read Archive (SRA). These transcriptome sequences were obtained from RNA-Seq experiments using Illumina NGS technology. After filtering by FASTX toolkit, the high quality clean reads were used for *de novo* assembly by Trinity using uniform parameters. Collectively, all cleaned high quality sequence read datasets contained 102.9 million reads, with more than 6.5 billion nucleotides (Table 1).

**Table 1:** Statistics of RNA-Seq experiments of mango cultivars.

| RNA-Seq<br>read datasets | References | Number of reads before<br>filtering | Number of reads after<br>filtering | Read length | Total bases |
| --- | --- | --- | --- | --- | --- |
| *cv. Langra* | Azim et al. 2014 | 12,153,196 | 5,314,486 | 80 | 425,158,880 |
| *cv. Zill* | Wu et al. 2014 | 68,419,722 | 44,578,030 | 80 | 3,566,242,400 |
| *cv. Shelly* | Luria et al. 2014 | 83,251,214 | 38,820,608 | 90 | 1,552,572,180 |
| *cv. Kent* | Dautt-Castro 2015 | 22,018,116 | 14,210,006 | 72 | 1,023,120,432 |

The four RNA-Seq datasets were assembled individually resulting in four datasets of unigenes (Table 2). The N50 of the assembled transcripts datasets were in the range of 525 – 1598 nucleotides (Table 2). The number of unigenes were as follows; cv. *Langra* = 30,953; cv. *Zill* = 58,797; cv. *Shelly* = 57,544; and cv. *Kent* = 85,036. The number of unigenes obtained as Trinity outputs were comparable as previously reported in respective publications (Azim et al. 2014; Wu et al. 2014; Luria et al. 2014; Dautt-Castro et al. 2014). This observation provided confidence for comparative transcriptome analysis.

**Table 2.** Statistics of unigenes data of mango cultivars assembled by Trinity program in this study.

| RNA–Seq read<br>datasets | N50 of unigenes<br>(bases) | Total length of<br>unigenes<br>(bases) | Total number of unigenes | Length of largest<br>unigenes (bases) | No. of unigenes reported<br>in the respective paper |
| --- | --- | --- | --- | --- | --- |
| *cv. Langra* | 525 | 14,216,135 | 30,953 | 5,821 | 30,509 (Azim et al. 2014) |
| *cv. Zill* | 1,050 | 40,388,175 | 58,797 | 11,243 | 54,207 (Wu et al. 2014) |
| *cv. Shelly* | 1,598 | 49,681,799 | 57,544 | 8,723 | 57,544 (Luria et al. 2014) |
| *cv. Kent* | 1,017 | 58,412,829 | 85,036 | 8,305 | 80,969 (Dautt-Castro et al.<br>2015) |

## Functional Annotation of Assembled Unigenes

To characterize the putative functions of mango unigenes, three different BLAST sequence similarity search strategies were adopted.

**(1)** All unigenes datasets were aligned against NT (non-redundant nucleotide sequence database), NR (non-redundant translated sequence database), Plant NT/NR, SwissProt and UniProt databases using BLAST. Sequence similarity searching showed homology of 83-96% unigenes with sequences in Nt and Nr databases.

**(2)** The unigene sequences were also compared with genomic sequence datasets of different Viridiplantae which revealed considerable sequence similarity with *Cirus sinensus*, *Citrus clementina*, *Populus tricarpa*, *Vitis vinifera* and *Riccinus communis* (Figure 1).

**Figure 1:**
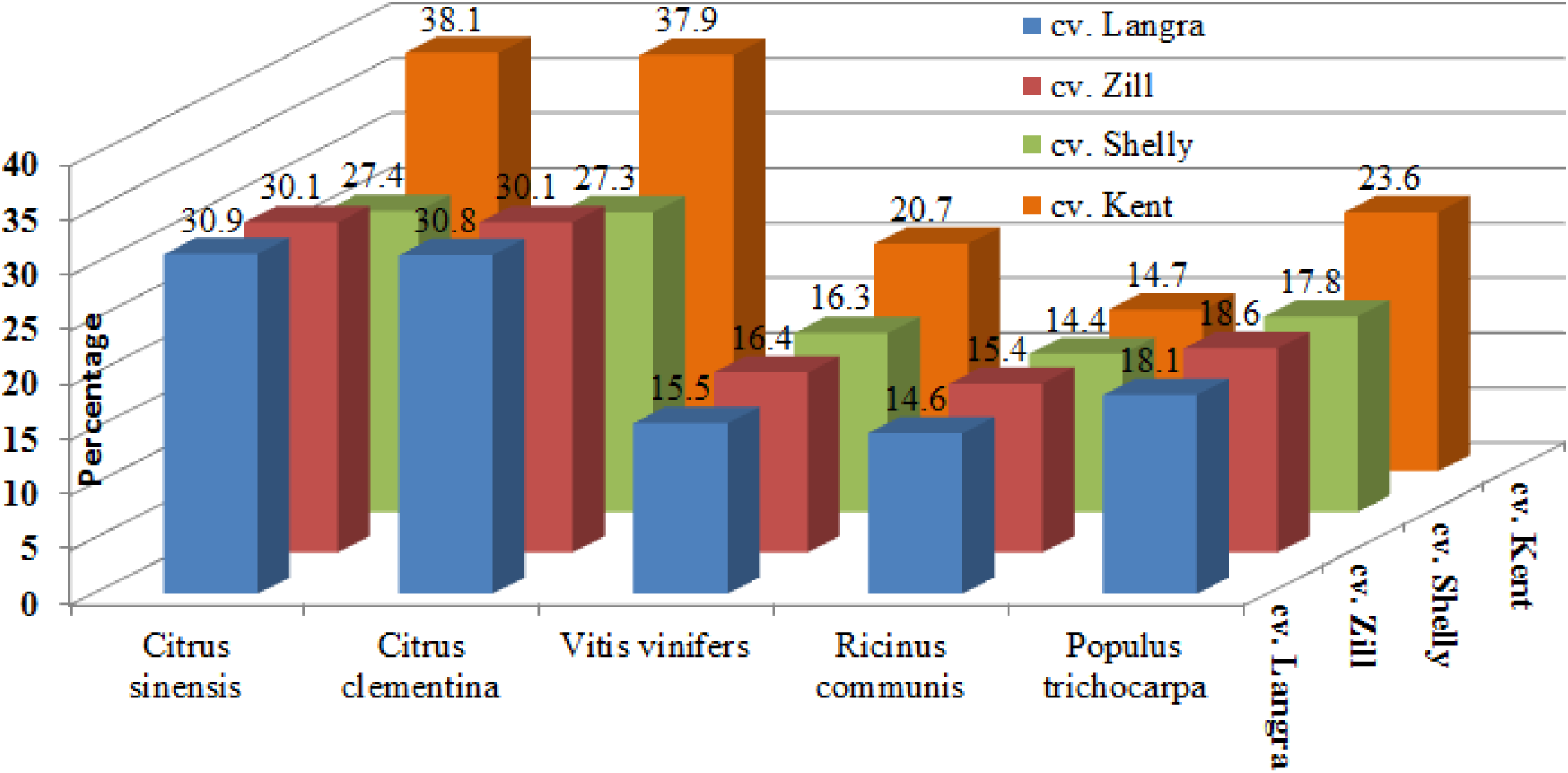
Comparative species distribution of mango cultivars transcriptomes.

**(3).** To find out cultivar-specific and ‘consensus’ sequences (i.e. unigenes common in four mango datasets studied); unigenes of four cultivars were compared to each other using BLAST. At every instance, one unigenes dataset was selected as query while other three datasets were considered as database. BLAST searches showed that 80–98% unigenes sequences of four mango cultivars have ≥75% identity with each other.

InterProScan provides a systematic language to describe the attributes of genes and gene products, which includes the functional characterization and annotation in combination with different protein signature recognition methods into one resource (Jones et al. 2014). InterProScan identified 12,388, 29,303, 25,878, and 18,793 protein coding sequences in cv. *Langra*, cv. *Zill*, cv. *Shelly* and cv. *Kent* unigenes datasets respectively. Functional characterization by Gene Ontology (GO) annotated an array of expressed genes in these mango varieties. GO analysis identified 17,704 (cv. *Langra)*, 18,846 (cv. *Zill*), 18,325 (cv. *Shelly*) and 12,119 (cv. *Kent*) GO terms, assigned to one of the three biological domains (i.e. Biological, Cellular and Molecular functions).

### Genes Related to Mango Flavor

Flavor of mango i.e. taste and aroma is constituted by a complex mixture of natural products. More than 500 volatile compounds have been reported to contribute in mango aroma and taste (Singh et al. 2013). The mango cultivars compared during present study have characteristic flavor (taste and aroma), color and consistency of pulp which is supposed to be due to different terpenoids, flavonoids and carotenoids biosynthetic pathways. Analysis of RNA-seq datasets of mango cultivars revealed unigenes involved in biosynthesis of oxygenated volatile compounds including esters, furanones and lactones. These secondary metabolites contribute as determinants of the characteristic aroma (Quijano et al. 2007). Amount and type of volatile compounds in mango often depend on area of production. Asian mangoes have more oxygenated volatile compounds such as esters, furanones, and lactones which give pineapple- or peach-like aromas to some varieties (Moshonas and Shaw 1994), while western mangoes that are hybrids of Asian stock have higher levels of certain hydrocarbons (Moshonas and Shaw 1994; MacLeod and de Troconis 1982). KEGG pathway analysis of four mango unigenes datasets identified active biochemical pathways involved in mango flavor, color and antioxidant activity.

#### Terpenoids

Terpene hydrocarbons are considered to be important contributors to flavor in most of the mango varieties (Quijano et al. 2007; El-Hadi et al. 2013). Many monoterpenes (C10) and sesquiterpenes (C20) comprise the most abundant group of compounds present in the aroma profile (Lichtenthaler 1999). The terpenoids are synthesized from two universal precursors isopentenyldiphosphate (IPP) and dimethylallyldiphosphate (DMAPP) which are products of two independent pathways: the cytosolic mevalonate (MVA) pathway; and plastidic1-deoxy-dxylulose-5-phosphate (DOXP) pathway (Nagegowda 2010). IPP and DMAPP are metabolized by a series of synthases (FDPS; farnesyldiphosphate synthase [EC 2.5.1.1], FPPS; farnesyldiphosphate synthase [EC 2.5.1.10] and GGPS1; geranylgeranyldiphosphate synthase, type III [EC 2.5.1.29]) into Geranyldiphosphate (GPP), Geranylgeranyldiphosphate (GGPP), and Farnesyldiphosphate (FPP). GPP processed to form monoterpenoids, whereas FPP enters in sesquiterpenoid/triterpenoid, carotenoid and N-glycan biosynthesis pathways (Figure 2). The analysis of unigenes of mango cultivars showed that GGPP acts as precursor of phytyldiphospahte (PyPP) and nona-prenyldiphosphate (NoPP) by all-trans-nonaprenyldiphosphate synthases [EC 2.5.1.85; EC 2.5.1.82]. PyPP and NoPP enter in Ubiquinone and other terpenoid-quinone biosynthesis pathways. Interestingly, in cv. *Shelly* dataset, two additional precursors i.e. hexaprenyldiphosphate and decaprenyldiphosphate are formed as indicated by unigenes encoding hexaprenyldiphosphate synthase [EC 2.5.1.83] and decaprenyldiphosphate synthase [EC 2.5.1.91]. These cv. *Shelly* specific precursors enter in Ubiquinone and other terpenoid-quinone biosynthesis pathway (Figure 2).

**Figure 2:**
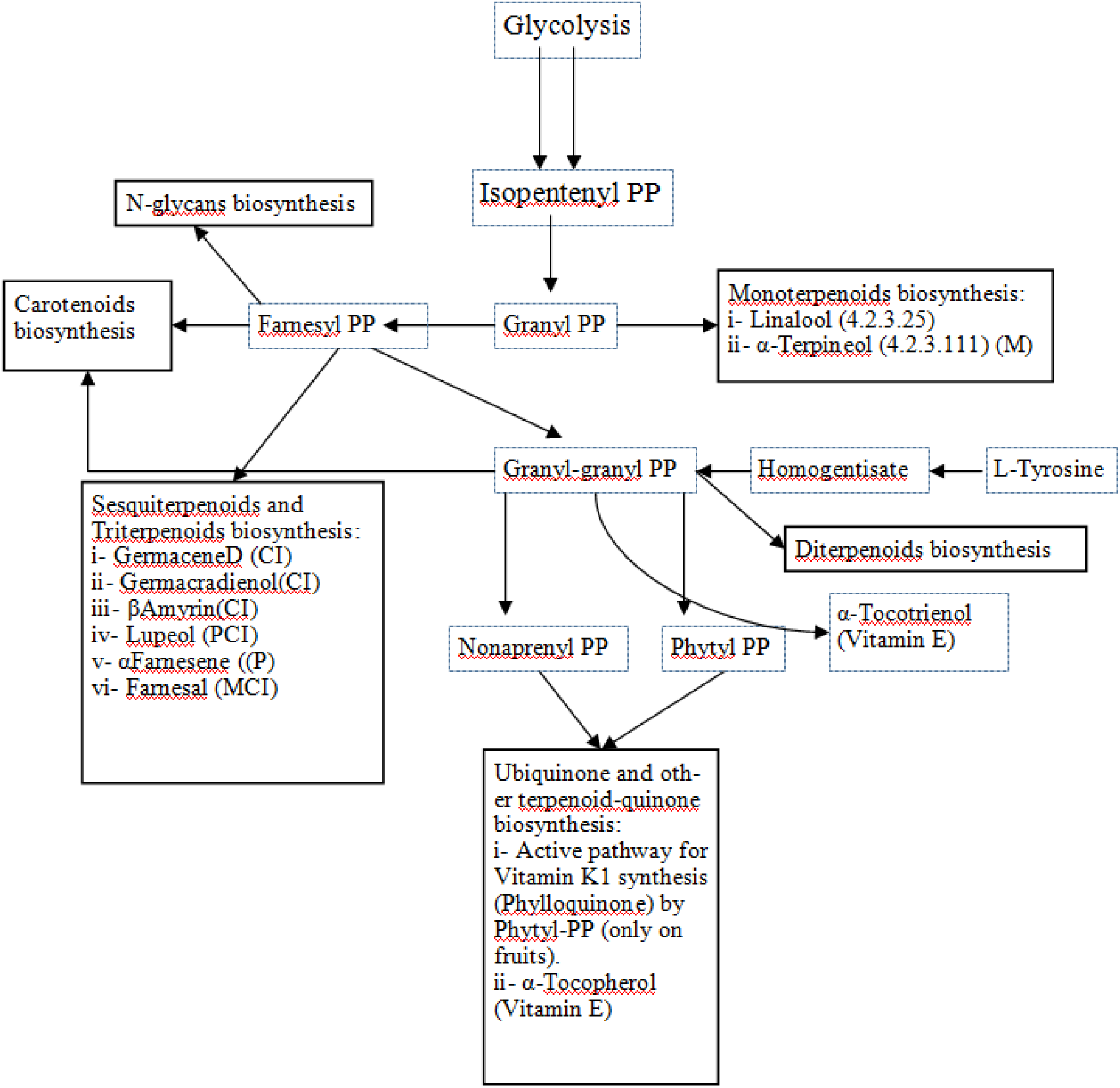
Terpenoid backbone biosynthesis in mango cultivars.

#### Monoterpenoids

Linalool and α-Terpineol are monoterpene alcohols, mainly found in flowers and spice plants having important commercial applications. These alcoholic compounds are responsible for pleasant aroma in fruits (Pino and Mesa 2006). The unigenes encoding SLinalool synthase [EC 4.2.3.25] responsible for synthesis of Linalool was found in all mango cultivars studied (Figure 2). This observation indicated that Linalool is the main monoterpene found in mango; while transcript encoding α-Terpineol synthase [EC 4.2.3.111] for α-Terpineol biosynthesis from Geranyl PP was found specifically in cv. *Shelly*. Among other monoterpenes, myrcene is found to be the major compound in most New World mango cultivars, along with sesquiterpene hydrocarbons which present in amounts as high as 10% (Lewinsohn et al. 2001).

#### Diterpenoids

Two diterpene biosynthetic pathways were found active in mango transcriptome datasets. (i) We found unigenes encoding enzymes involved in Gibberellins hormone biosynthesis produced from geranylgeranyldiphosphate (GGPP) via ent-copalyldiphosphate by the bifunctionalent-copalyldiphosphate/ent-kaurene synthase (CPS/KS). Gibberellins are tetracyclic diterpenes (C20) which stimulate wide variety of responses during plant growth (Phinney and Spray 1987). (ii) A unigene encoding homoterpene synthase [EC 1.14.13.B14] (a cytochrome P450 enzyme) was also found except in cv. *Kent* dataset. This enzyme catalyses the conversion of secondary metabolite geranyl linalool to the homoterpene 4,8,12-trimethyltrideca-1,3,7,11-tetraene (TMTT) (C16). TMTT is an insect-induced volatile compound involved in plant defense response. Terpene volatiles play a vital role in plant-organism interactions as attractants of pollinators or as defense compounds against herbivores (Lee et al. 2010).

#### Sesquiterpene and Triterpenoids

A number of unigenes encoding enzymes responsible for biosynthesis of sesquiterpenes and triterpenes were found in mango transcriptome datasets (Figure 2). (i) The unigenes encoding germacradienol synthase [EC 4.2.3.22] which catalyzes the sesquiterpenoid Germacrene D biosynthesis, was found in datasets of cv. *Zill* and cv. *Shelly*. This sesquiterpene is reported to have antimicrobial and anti-insecticidal activities (Noge and Becerra 2009). (ii) The unigenes encoding lupeol synthases [EC 5.4.99.-] and β-amyrin synthase [EC 5.4.99.39] for biosynthesis of pharmacologically active triterpenoids, i.e. Lupeol and β-amyrin were found in cv. *Langra*, cv. *Zill* and cv. *Shelly* datasets (Saleem 2009; Siddique and Saleem 2011; Santos et al. 2012).

Besides terpenoids, other compounds also contribute in mango flavor and fragrance. Mango possesses a very attractive sweet essence characteristic due to the presence of different sugars. The major sugars found in mango are glucose, fructose, and sucrose (MacLeod and de Troconis 1982). In accordance with the literature, unigenes encoding enzymes responsible for the synthesis of sucrose, fructose and glucose were found in all four cultivars. In addition, the cv. *Langra* dataset have transcripts encoding enzymes of mannose and galactose pathways; whereas 17 transcripts encoding enzymes of pentose sugar pathway were detected in cv. *Zill* and cv. *Shelly* datasets.

Among the carbonyls, 14 unigenes encoding series of oxidoreductases for biosynthesis of (E)-2-hexenal and hexanal were found in fruit transcriptome datasets (i.e. cv. *Kent*, cv. *Shelly* and cv. *Zill*) while absent in leaf dataset (cv. *Langra*). These compounds have fatty–grassy and green–fruity notes that could make a minor contribution to mango aroma and reported to be bactericidal (Pino and Mesa 2006; Lanciotti et al. 2003). Several unigenes encoding esterases which hydrolyse fruit lactone as intramolecular esters of 4- and 5-hydroxy acids were also found. These were previously characterized in mangos (Moshonas and Shaw 1994; Macleod and de Troconis 1982), and are considered to be important contributors to the flavor and aroma of this fruit (Pino et al. 2005; Fahlbusch et al. 2007).

### Genes Involved in Carotenoids and Flavonoids Biosynthesis

Plant carotenoids are tetraterpenes and the most vital colored phytochemicals which occurs as all-trans and cis-isomers (Khoo et al. 2011). This group of natural products referred as pigment and nutraceuticals (Botella-Pavía et al. 2004) accounting for the brilliant colors in fruits and vegetables (Khoo et al. 2011). Carotenoids also act as a precursor for the production of apocarotenoid hormones such as abscisic acid which regulate development of plant and its interaction with their environment (Nambara and Marion-Poll 2005). Carotenoids are derived from the 40-carbon isoprenoid phytoene that participate in light harvesting and essential for photoprotection against excess light (Ruiz-Sola and Rodríguez-Concepción 2012; Moran and Jarvik 2010). Plant carotenoids are red, orange, and yellow lipid-soluble color pigments embedded in the membranes of chloroplasts and chromoplasts (Bartley and Scolnik 1995). The most studied carotenoids include β-carotene, lycopene, lutein and zeaxanthin. However, the intensity of color in fruits and vegetables depends on the concentration of carotenoids and their growth maturity (Khoo et al. 2011).

Lycopene and β-carotene are important carotenoids of mango (Khoo et al. 2011). Unigenes encoding enzymes for lycopene and β-carotene were found in cv. *Zill* and cv. *Shelly* fruits datasets giving a reddish-orange color to the fruit (Figure 3) (Wu et al. 2014; Luria et al. 2014). On the other hand, absence of β-carotene and presence of lycopene in cv. *Kent* is predicted to be responsible for orange-yellow color of the fruit (Dautt-Castro et al. 2015).

**Figure 3:**
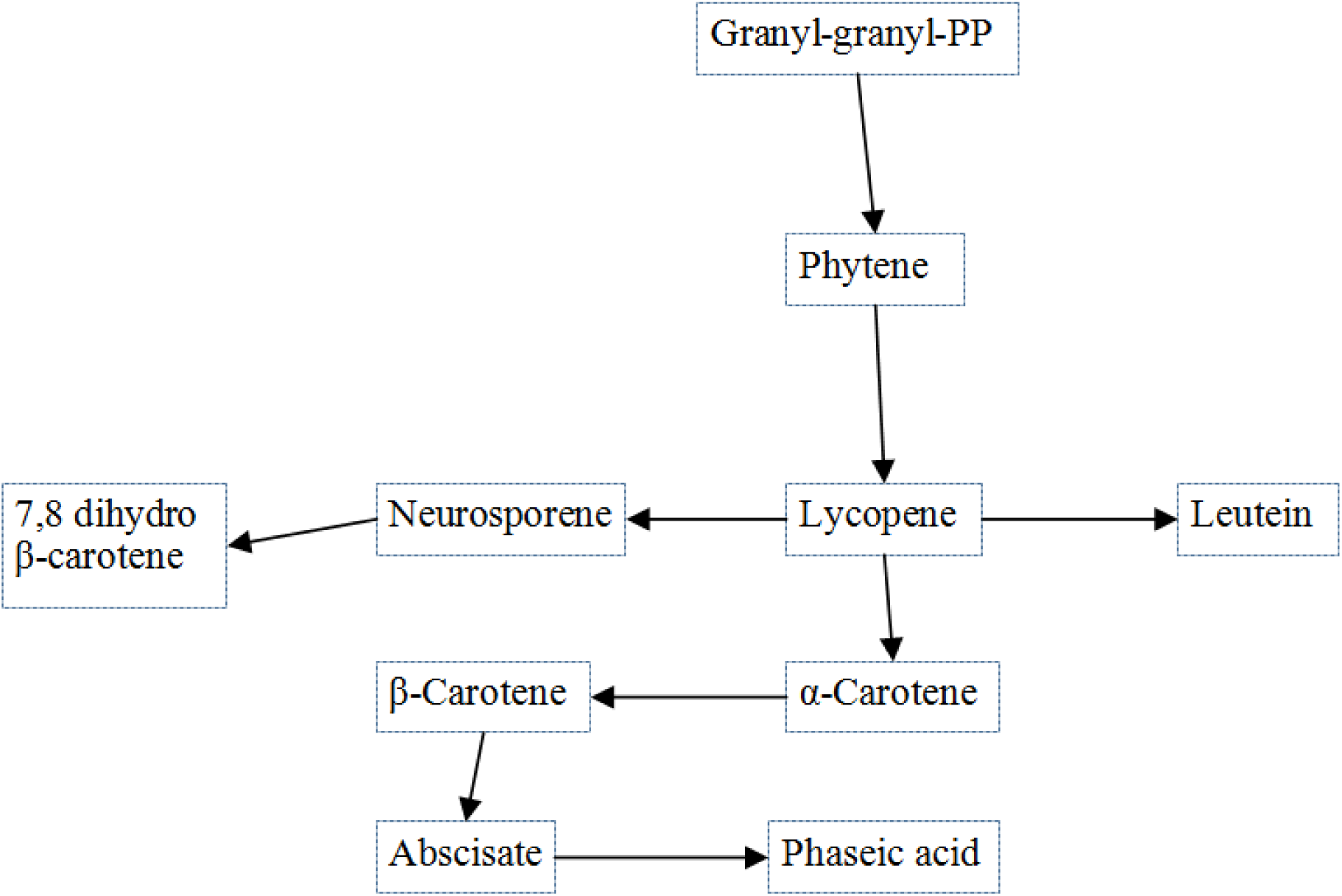
Carotenoid biosynthesis in mango cultivars.

Many unigenes encoding enzymes and related proteins involved in biosynthesis of flavonoids, the group of pigments that color most flowers, fruits, and seeds were present in four mango cultivars datasets (Figure 4). These flavonoids included naringenin, pinobanksin, afzelechin, apiferol, eriodictoyl, luteolin, catechins (epicatechin, gallocatechin, epigallocatechin) myricetin, and quercetin. Flavonoids are phenylpropanoid-derived plant metabolites and ubiquitous in nature (Hoang et al. 2015). According to chemical structure, these secondary metabolites are classified into flavonols, flavones, flavanones, isoflavones, catechins, anthocyanidins and chalcones. Flavonoids are known to perform diverse functions including color-based attractants to pollinators and symbionts (Dixon and Pasinetti 2012). In higher order plants, flavonoids are also required for UV filtration, nitrogen fixation, cell cycle inhibition, and as chemical messengers. These compounds also act as allelochemicals, antimicrobial, antiherbivore, antiallergic, antiplatelet, anti-inflammatory, anti-tumor and antioxidant agents (Falcone Ferreyra et al. 2012).

**Figure 4:**
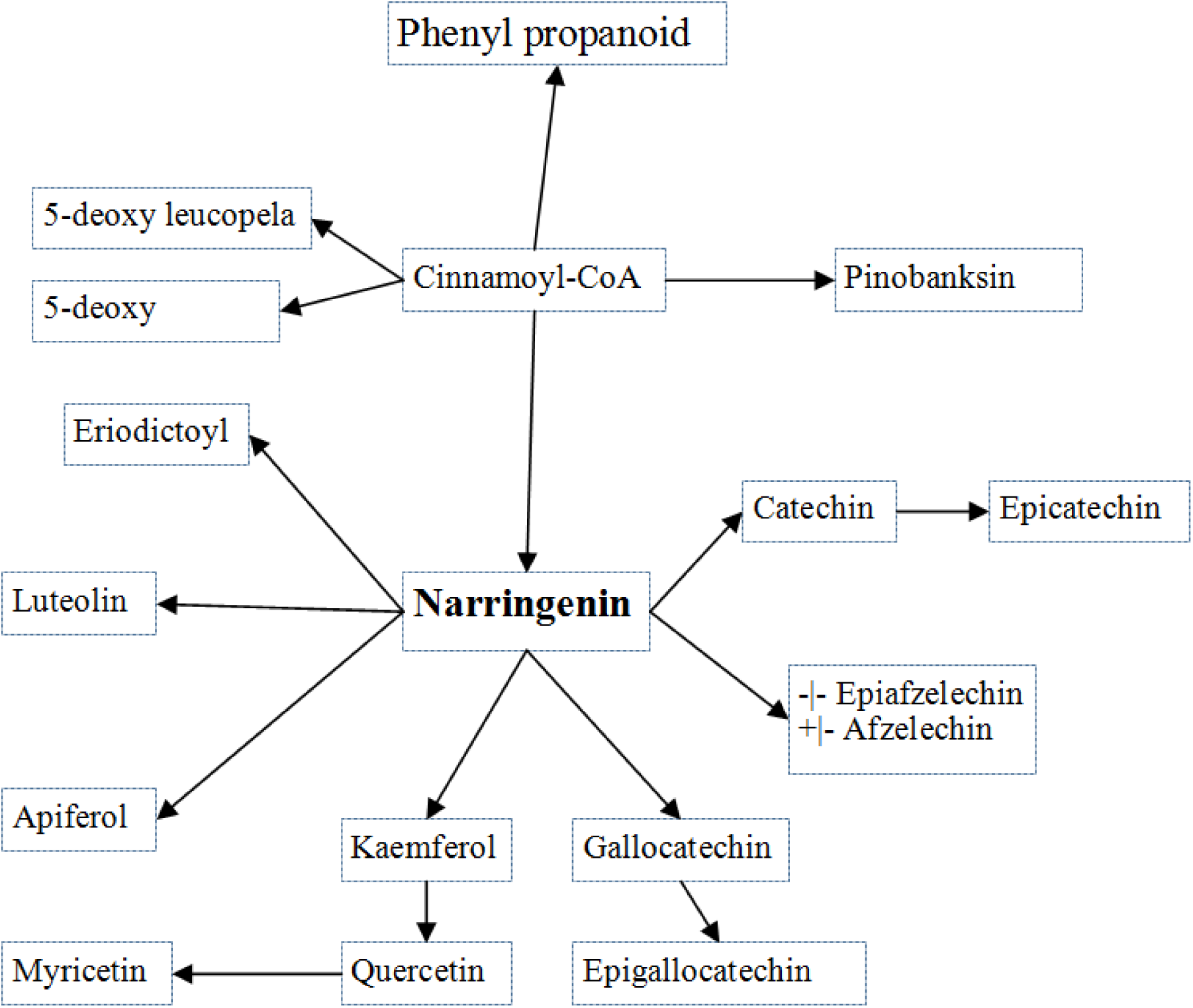
Flavonoid biosynthesis in mango cultivars.

### Genes involved in biosynthesis of antioxidants

Tocopherols and tocotrienols (vitamin E), ascorbic acid (vitamin C) and carotenoids react with free radicals and reactive oxygen species, which is the basis for their function as antioxidants (Sies and Stahl 1995). The presence of active pathways for biosynthesis of Vitamin E and C and carotenoids in all four cultivars of mango signifies presence of the antioxidant activity.

The present comparative transcriptomic analysis of four mango cultivars from different countries provided insight of genes encoding for enzymes for biosynthesis of terpenoids, carotenoids, flavonoids and other natural products. These volatile and nonvolatile metabolites are determinants of flavor (aroma and taste) and color.

